# Comparison of hybrid learning and remote education in the implementation of the “Adopt a Microorganism” methodology

**DOI:** 10.1101/2021.03.09.434548

**Authors:** Bárbara Rodrigues Cintra Armellini, Alexandre La Luna, Vanessa Bueris, Alisson Pinto de Almeida, Alicia Moraes Tamais, Flávio Krzyzanowski Júnior, Victor Samuel Hasten Reiter, Camilo Lellis-Santos, Rita de Cássia Café Ferreira

**Affiliations:** Institute of Biomedical Sciences, University of São Paulo, São Paulo, Brazil; Federal Institute of São Paulo, Campus Sorocaba, São Paulo, Brazil; Federal Institute of São Paulo, Campus Capital, São Paulo, Brazil; Federal University of São Paulo, Campus Diadema, São Paulo, Brazil

## Abstract

The Internet has changed the way teachers and students access information and build knowledge. The recent COVID-19 pandemic has created challenges for both teachers and students and a demand for new methodologies of remote learning. In the life sciences, mixing online content with practical activities represents an even greater challenge. In microbiology, the implementation of an active teaching methodology, the #Adopt project, based on the social network Facebook^®^, represents an excellent option for connecting remote education with classroom activities. In 2020, the version applied in high school, “Adopt a Microorganism”, was adapted to meet the demands of emergency remote education owing to the suppression of face-to-face activities caused by the pandemic. In the present study, we assessed how the change in methodology impacted the discourse richness of students from high school integrated with technical education in the Business Administration program of the Federal Institute of São Paulo, Sorocaba Campus. Three questionnaires related to the groups of microorganisms (Archaea, Bacteria, Virus, Fungi, and Protozoan) were applied. The students’ responses in the 2019 and 2020 classes were compared concerning content richness and multiplicity of concepts through the application of the Shannon diversity index, an approach that is generally used to assess biodiversity in different environments. The observed results suggest that remote learning provided students with a conceptual basis and richness of content equivalent to that achieved by students subjected to the hybrid teaching model. In conclusion, this study suggests that the #Adopt project methodology increases students’ discourse richness in microbiology even without face-to-face traditional classes.

## INTRODUCTION

Internet access is growing daily, with Brazil in a prominent position, as together with India and China, it is responsible for about 70% of access, ranking 4th in this topic (1). Additionally, approximately 70% of the Brazilian population is connected to the Internet (2), with social media occupying an important space. According to the “We Are Social” website (3), 62% of Brazilians are connected to social networks, especially Facebook^®^, which has just over 127 million users in Brazil, according to data from the website.

These facts show the relevance of the Internet in our lives and, consequently, in the lives of students, who now have immediate and unrestricted access to content that would previously have only been found in textbooks or transmitted by the teacher (4). This scenario makes it even more difficult to apply a traditional teaching model, in which the central role is that of the teachers (5), as they are no longer the sole holders of knowledge. Therefore, it is interesting to think about the Internet as an ally in pedagogical practices, as it promotes an expansion of learning environments and diversifies mobilization strategies (6).

The Internet allows students to take a more active role in their learning, which facilitates the connection of learned concepts and promotes the development of critical thinking (7). Moreover, the placement of the student as a central figure in the development of their activities makes teaching more meaningful and thus creates the need to seek concepts that allow for a greater understanding of a subject or idea that awaken the student’s interest in learning (8).

Among all the possibilities of Internet use, social media is significant because of its relevance and wide reach, as previously mentioned. Several studies have shown that its use has contributed in some way to learning, both in higher education and in high school (9; 10). Among all the existing social networks, one of the most used by teachers and students is Facebook^®^ (11). Santos and Campos (2014) (12) demonstrated that its use for educational purposes is compelling, as discussions and teacher-student relationships can be expanded, compared to traditional teaching methods.

In 2013, the #Adopt Project was created using Facebook^®^ as a didactic platform for teaching microbiology, both in higher education (“Adopt a Bacterium” (13)) and in high school (“Adopt a Microorganism”). The latter was applied to students in the 2nd year of high school integrated with technical education in Business Administration of the Federal Institute of São Paulo, Sorocaba Campus, during 2019 and 2020.

In 2019, the “Adopt a Microorganism” project was applied in its original hybrid version, with alternating face-to-face classes and practice with online activities on Facebook^®^. In 2020, however, education experienced drastic changes with the emergence of the COVID-19 pandemic. Schools and colleges were closed, and, in most cases, classes began to take place online without prior planning or training (14), forcing both teachers and students to adapt to a new system in a very short time. This project had just started before the pandemic lockdown and had to be adapted to be offered completely online.

Given these differences, we assessed how the change in the “Adopt a Microorganism” methodology impacted the gain of terminology in microbiology by high school students. The acquisition of biology terminology is an essential skill during biology learning (15), and we measured the increase of this vocabulary through the richness of speech. For this purpose, we used the Shannon diversity index (16), which is widely employed in conservation biology (17; 18) and in microbiology to assess the richness of microbial populations (19; 20). In addition, we analyzed the perceptions of students regarding the strengths and weaknesses of the project.

## MATERIALS AND METHODS

### Study design

This study, in both 2019 and 2020 editions, was conducted with high school students from the 2nd year of the Federal Institute of Education, Science, and Technology of São Paulo, Sorocaba Campus. The “Adopt a Microorganism” project was applied during the first semester with 36 students in 2019 and 38 in 2020. The two cohorts were similar, especially because they have entered the school through the same selection process, have similar socioeconomic status, and belong to the same age group. There was also gender and ethnic equivalence. The students were distributed into five groups, each responsible for a group of microorganisms: Archaea, Bacteria, Virus, Fungi, or Protozoan. In addition to the lectures regarding basic microbiology topics, the teacher posted weekly challenges (S1 Appendix) on Facebook^®^, based on the PISA (International Student Assessment Program) and ENEM (National High School Exam), with increasing degrees of difficulty (easy, medium, and difficult) using Bloom’s taxonomy (21). The first three weeks included the same general challenges for all groups regarding the biological definition of life and the main differences between the three domains of life. In the last three weeks, the challenges were specific to each group and promoted discussion about the “adopted” microorganism, its relationship with humans, and the diseases caused by it. Student responses and discussions on Facebook^®^ were mediated by undergraduate or graduate students who had already participated in the project and received a brief training (22).

The mediators were different for each group. Periodic meetings were held with the mediators. They were instructed to check the reliability of the sources used by the students, to point out possible conceptual errors, and not to provide the students with ready answers, but rather encourage them to research new information to supplement or correct their answers. The class teacher, who was the same in both years, supervised all stages of the “Adopt a Microorganism” and also the mediators’ performance, to ensure that all groups made similar progress throughout the activities. Each challenge needed to be validated by the mediators, when they thought that the objective had been reached, to compute the student’s score.

At the end of all the activities, the students produced promotional material aimed at the lay public, to provide society with scientific information. In 2019, the students presented a seminar on their adopted microorganisms to their classmates and other school members. They also participated in hands-on practical activities (“Journey into the world of microorganisms and human parasites”) that were held immediately after the end of the general challenges on Facebook^®^ at São Paulo University. In these activities, the students could reinforce their knowledge about their adopted microorganisms through several laboratory activities, such as seeding and cultivation techniques, microscopic analysis of microbiologic material, technical visits to research laboratories, and participation in ludic games alongside the mediators. Because of the COVID-19 pandemic, face-to-face classes were interrupted in 2020, and the students could not participate in the hands-on practical activities or present seminars. Therefore, they prepared mental maps as promotional material concerning the adopted microorganisms. During a semester, the Facebook discussions lasted six weeks and the students had another two weeks to prepare the seminars/promotional material.

### Data collection

We administered three voluntary and anonymous surveys to evaluate the students’ knowledge of their adopted microorganisms. The students answered the question, “What do you know about the adopted microorganism?” The surveys had an open question, and a minimum or a maximum number of lines that could be written by students was not stipulated. In addition, they were instructed to write down everything they knew/remembered about the studied microorganism, without consulting and without worrying about grading, as the answers were anonymous. All students answered the questionnaires at the same time. The first survey (Q1) was conducted before the start of the discussions on Facebook^®^. In the 2019 edition, the second survey (Q2) was conducted after the hands-on practical activities, and in 2020, it was applied immediately after the end of the discussions on Facebook^®^. The last survey (Q3) was conducted five months after the second survey was administered, to evaluate the students’ knowledge retention. Plagiarism was checked by the master’s student responsible for analyzing the results and by the class teacher, both in the students’ answers to the questionnaires and in the answers to the Facebook challenges. In the latter, the mediators also checked for plagiarism.

To assess the feasibility of using Facebook^®^ as a teaching and learning platform for microbiology in biology classes, an anonymous and non-mandatory questionnaire was administered to the participating students at the end of the project (S2 Appendix). This survey contained questions such as “point out the positive and negative aspects of using Facebook^®^ for learning microbiology”. In the 2020 edition, because of the suspension of classes due to the pandemic and the continuity of the project in an exclusively remote format, two questions were included, one to determine whether the continuity of the project allowed for communication and contact with colleagues and another to determine whether it was possible to continue learning and studying at home. Thirty-two students responded, and these responses were analyzed and categorized based on the extraction of the nuclei of meaning according to Bardin (1977) (23).

### Data analysis

For each survey, the students’ responses were evaluated to identify the presence and quantity of each of the following topics: taxonomy, morphology, metabolism, genetics, pathogenicity, treatment, prevention, ecology, reproduction, life cycle, social impact examples, and others. For further details on the criteria to classify the students’ phrases in the questionnaires, see Table S1. In addition, we evaluated whether the topic discussed in the responses had any conceptual errors. Subsequently, for each topic, we computed the percentage of students that commented on the topic at least once. For this, we have used a custom Python 3.8 script, following the equation:

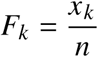

where *F*_*k*_ is the number of students that commented at least one time about the topic k (*x*_*k*_) divided by the number of students in the class (*n*). The same was computed for the conceptual errors on each topic.

To assess the average diversity of topics worked on by each group in each questionnaire, we calculated the Shannon diversity index (H) for each group (16). This index is not widely used to evaluate teaching, but it can be an interesting approach when the data obtained involve specific knowledge categories and the frequency with which these categories are present in each speech given or written by students.

For this calculation, we used the ecopy 0.1.2.2 library for Python 3.8, following the equation:

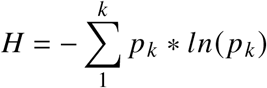

where *p*_*k*_ is the frequency of appearance of each topic in the responses of each group, and k is the total number of different subjects that appear in the answers. Thus, for each questionnaire, we obtained the Shannon index per group for the whole class and conceptual errors. The higher this index, the greater the diversity of the topics covered in the responses. The confidence intervals for each index were estimated by bootstrapping, as in Laura Pla (2004) (24). To compare Shannon’s indices between each questionnaire, we performed a univariate analysis of variance (ANOVA), after testing for homogeneity of variance (Levine’s test). As this analysis was performed for each year, the significance level in the univariate ANOVAs was adjusted downward by a Bonferroni’s correction (α = 0.025). The ANOVAs were followed by two-tailed t-tests corrected for multiple comparisons also employing Bonferroni’s correction (α = 0.004). The statistical significance analysis was conducted using jamovi 1.6 software. Complementary to these analyses, we calculated the difference between the indices (effect size) and estimated the 95% confidence interval for the difference(26).

Furthermore, to assess the degree of dissimilarity between Q1 and Q2 and between Q2 and Q3 for each year and between the 2019 and 2020 editions, we calculated the Bray-Curtis dissimilarity between these questionnaires (27). To calculate the Bray-Curtis dissimilarity, we used the ecopy 0.1.2.2 library for Python 3.8, following the equation:

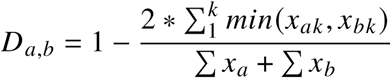

where k is the total number of topics that appeared in questionnaires “a” and “b”, and x_*ak*_ and x_*bk*_ are the number of appearances of topic k in questionnaires “a” and “b”, respectively. The greater the Bray-Curtis dissimilarity, the larger the difference between the compared questionnaires. The confidence intervals for each dissimilarity were estimated by bootstrapping, as in Laura Pla (2004) (24).

Lastly, we also counted the number of words in the questionnaire answers using the NLTK 3.5 library for Python 3.8. To measure the correlation between the number of words and the number of errors in Q1, Q2, and Q3, we calculated Pearson’s r and its 95% CI. For this, we used the pingouin 0.3.8 package for Python 3.8. Additionally, using Python 3.8, with the statsmodels 0.12.1 module, we built a linear regression model, relating the number of errors to the number of words used.

### Data Plot

For data visualization, we built word clouds for the answers to each survey by group. The clouds were built using a custom Python 3.8 script. First, the answers were preprocessed with a tokenization and lemmatization step using the NLTK 3.5 library. We also removed the punctuation and stopwords. In some specific cases, the words were joined together to avoid any loss of meaning in the student’s sentence (for example, “naked eye”). Word clouds were created with the WordCloud 1.8.1 library and edited with Matplotlib 3.3.2. The most prominent words were those that were most repeated in the students’ responses to the surveys.

To plot the graphs, we used another custom Python 3.8 script with the Seaborn 0.11.0 and Matplotlib 3.3.2 libraries.

### Ethics

This project (CEP ICB-USP Protocol #990/2018) was evaluated by the Research Ethics Committee of Human Beings (CEPSH ICB-USP) and was considered exempt from the need for consent form (report #1247 issued on 11/26/2018). This exemption was because the present study did not carry out any procedures regulated by CONEP resolution #466/2012.

### Limitations

In 2019, Q2 was applied shortly after the scientific hands-on practical activities held in the middle of the project (just after the end of the general challenges and before the specific challenges). In 2020, owing to the COVID-19 pandemic, these activities did not take place, and Q2 was applied only at the end of the project (after the specific challenges). Thus, we did not calculate the Bray-Curtis dissimilarity between the 2019 and 2020 Q2 questionnaires. In addition, in 2020, Q2 and Q3 were applied remotely (via Google Forms). Therefore, it was not possible to control whether the students used search functions for didactic materials and websites, which may have impacted the comparisons between 2019 and 2020 editions.

Lastly, as the mediators were not the same for all groups, and some changed from one year to another of the project, this may have resulted in small differences in the content worked on between different groups and different years, based on possible influences by mediators, both in engagement and content discussed with students. However, although there were no specific rubrics to guide the mediators throughout their activities, they met with the class teacher periodically to receive guidance on the different stages of the project.

## RESULTS

To assess the students’ knowledge about the adopted microorganisms before, during, and after the “Adopt a Microorganism” project implementation, three voluntary and anonymous questionnaires were administered to the high school students from the 2nd year of the Federal Institute of Education, Science, and Technology, Sorocaba Campus. In 2019, Q1 was applied just before the beginning of the project (and before microbiological classes took place), and 36 responses were obtained; Q2 was applied shortly after the hands-on practical activities with 34 responses; Q3 had 32 responses and was applied five months after the project ended. In 2020, Q1 and Q3 were applied at the same time point as in 2019, with 36 and 32 responses, respectively, and Q2 was applied at the end of the project, with 24 responses.

We observed a large number of short answers in the 2019 Q1, and they reflected, in most cases, common knowledge about the adopted microorganism. They generally addressed issues related to its pathogenicity or its social impact (benefits or harms produced by the microorganism) (Fig 1A). Conceptual errors, mainly related to the morphology and ecology of microorganisms (Fig 1C), were also common. This scenario changed in Q2, in which more students provided information about the adopted microorganism concerning its taxonomy, reproduction, and ecology (Fig 1A). Moreover, we observed fewer students committing conceptual errors and they focused mainly on the taxonomy and morphology (Fig 1C). In Q3, a higher percentage of students have written about the adopted microorganisms when compared to Q1, predominantly on reproduction, examples, and ecology (Fig 1A).

**Fig. 1.**
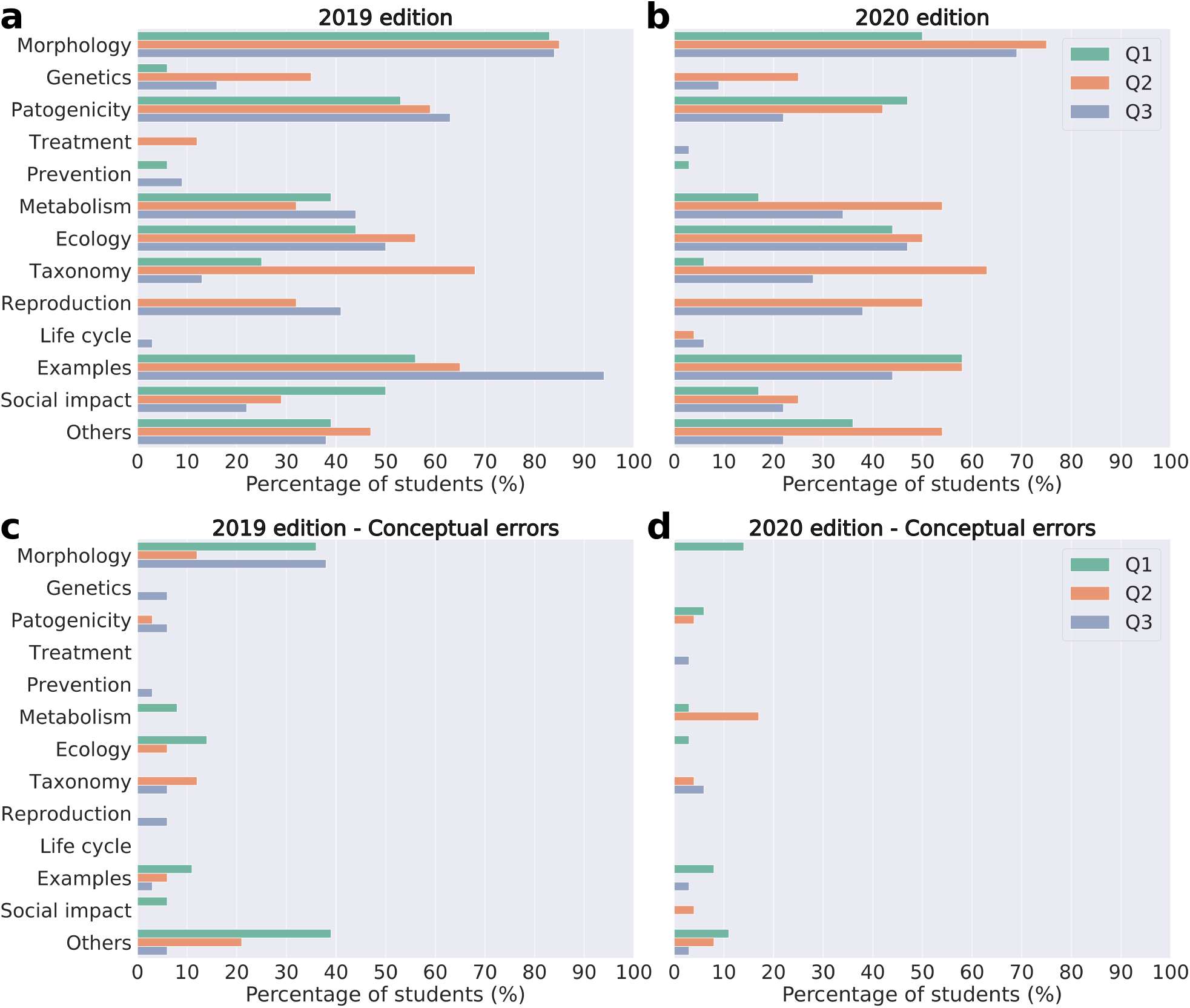
Percentage of students that commented on each topic. Percentage of students that commented on the topic at least once in the: Q1 survey, applied before the start of “Adopt a Microorganism”; Q2 survey, applied after the hands-on practical activities in 2019 and immediately after the end of the Facebook^®^ discussions in 2020; Q3 survey, applied five months after the end of the project. **[a]** microbiology topics that were cited correctly in 2019; **[b]** microbiology topics that were cited correctly in 2020; **[c]** microbiology topics that were wrongly cited (conceptual errors) in 2019; **[d]** microbiology topics that were wrongly cited (conceptual errors) in 2020.

Regarding the differences in the Shannon index, the ANOVA suggests that there is a difference between the Shannon indices of the Q1, Q2 and/or Q3 questionnaires (F(2, 96) = 4.582, p = 0.012, η_p_2 = 0.09) (Fig 2A). Considering all students, the content richness of the responses increased in Q2 compared to Q1 (H_Q2–Q1_=0.17, 95% CI=[0.07; 0.29], p < 0.001) (Fig 2A and Table 1). However, when comparing Q3 with Q1, the difference between Shannon index is lower and the confidence interval is also consistent with the absence of a difference, which is corroborated by the post hoc test (H_Q3–Q1_=0.08, 95% CI=[-0.05; 0.18], p = 0.020) (Fig 2A and Table 1). When comparing each group, a similar trend was observed, in which the effect size between Q2 and Q1 was, in general, larger than the effect size between Q3 and Q1 (Fig 2C and Table 1). Furthermore, these effect sizes and the confidence interval associated, except for protozoan and virus, remained predominantly positive. However, in most comparisons, the t-test and the confidence interval range do not rule out the possibility of no difference between the questionnaires (Fig 2C and Table 1).

**TABLE 1.**
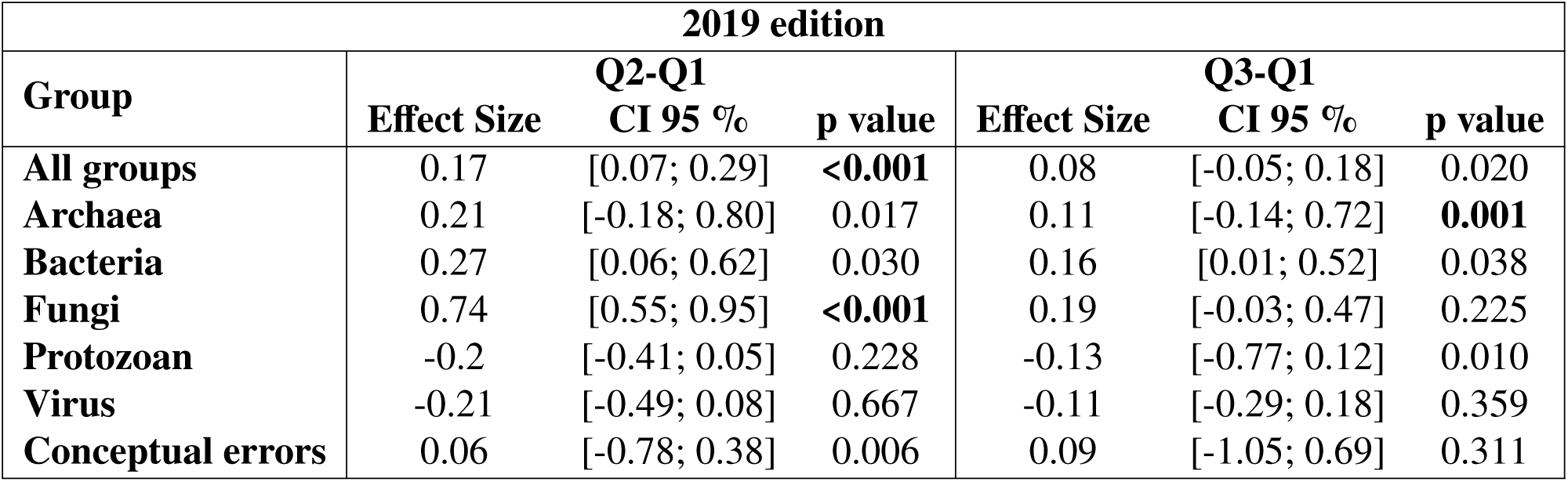
Differences in the Shannon index between the questionnaires Q1 and Q2, and Q1 and Q3 for the 2019 edition.

| **2019 edition** |  |  |  |  |  |  |
| --- | --- | --- | --- | --- | --- | --- |
| **Group** | **Q2-Q1** |  |  | **Q3-Q1** |  |  |
|  | **Effect Size** | **CI 95 %** | **p value** | **Effect Size** | **CI 95 %** | **p value** |
| **All groups** | 0.17 | [0.07; 0.29] | **<0.001** | 0.08 | [-0.05; 0.18] | 0.020 |
| **Archaea** | 0.21 | [-0.18; 0.80] | 0.017 | 0.11 | [-0.14; 0.72] | **0.001** |
| **Bacteria** | 0.27 | [0.06; 0.62] | 0.030 | 0.16 | [0.01; 0.52] | 0.038 |
| **Fungi** | 0.74 | [0.55; 0.95] | **<0.001** | 0.19 | [-0.03; 0.47] | 0.225 |
| **Protozoan** | -0.2 | [-0.41; 0.05] | 0.228 | -0.13 | [-0.77; 0.12] | 0.010 |
| **Virus** | -0.21 | [-0.49; 0.08] | 0.667 | -0.11 | [-0.29; 0.18] | 0.359 |
| **Conceptual errors** | 0.06 | [-0.78; 0.38] | 0.006 | 0.09 | [-1.05; 0.69] | 0.311 |

**Fig. 2.**
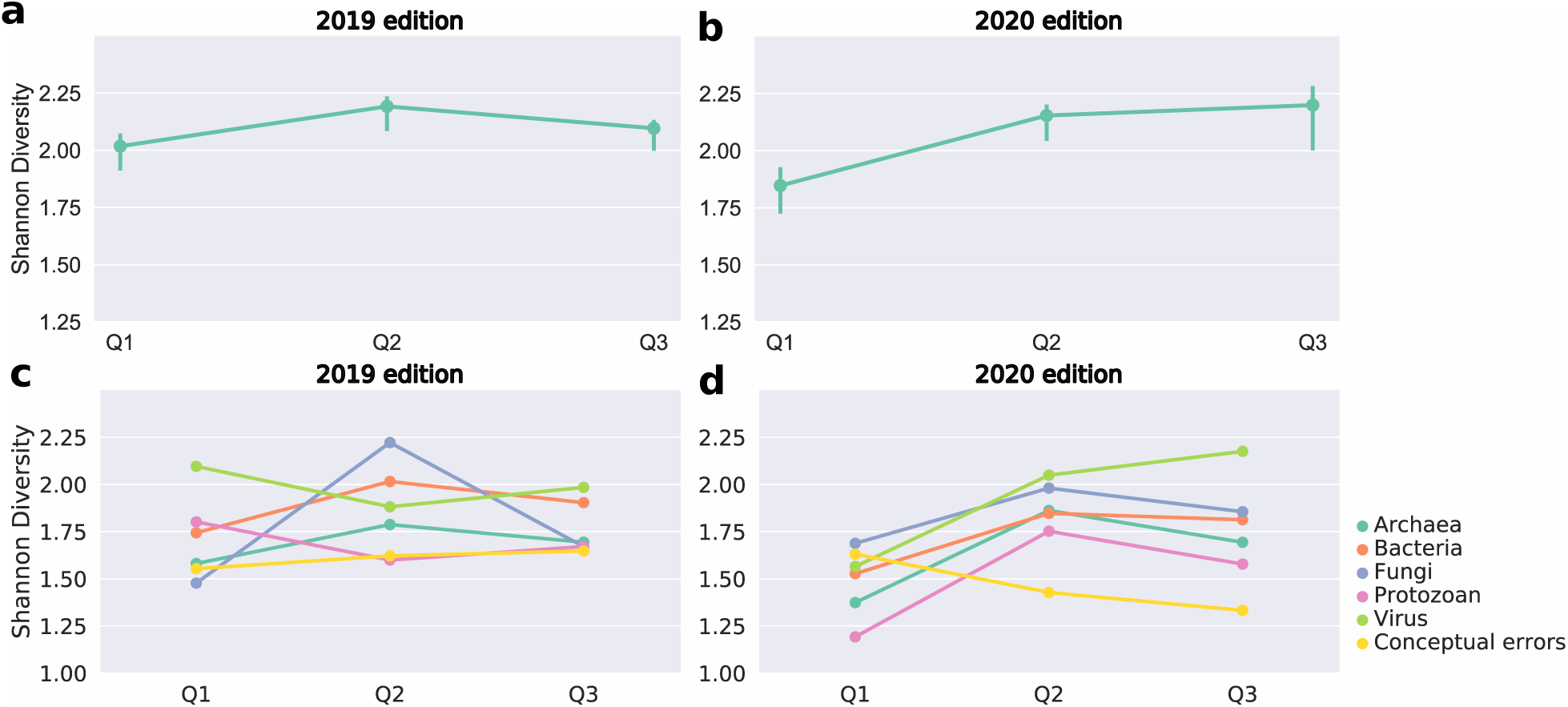
Word richness in the questionnaires. Graphs showing the variation in the student’s response richness (calculated by the Shannon index) over the three surveys: Q1, applied before the beginning of “Adopt a Microorganism”; Q2, applied after the hands-on practical activities in 2019 and immediately after the end of the Facebook^®^ discussions in 2020; Q3, applied five months after the end of the project. **[a]** variation in the Shannon index for all surveys analyzed in 2019; **[b]** variation in the Shannon index for all surveys analyzed in 2020; **[c]** variation in the Shannon index of the surveys answered by each group (Archaea, Bacteria, Fungi, Virus and Protozoan) in 2019; **[d]** variation in the Shannon index of the surveys answered by each group (Archaea, Bacteria, Fungi, Virus and Protozoan) in 2020.

This variation in the richness of the students’ responses between surveys was supported by the calculated Bray-Curtis dissimilarities (Table S2). In all cases, the dissimilarity was higher than 0.15, and the confidence interval was restricted, in most cases, to values greater than 0.15 and smaller than 0.50 (Table S2). The main exceptions were the virus group and conceptual errors (Virus: Q1–Q2=0.52, 95% CI=[0.45; 0.71]; Q2–Q3=0.44, 95% CI=[0.40; 0.61] | Conceptual errors: Q1–Q2=0.51, 95% CI=[0.51; 0.51]; Q2–Q3=0.55, 95% CI=[0.55; 0.55]). Moreover, it is interesting to note that the dissimilarity values between Q1–Q2 and Q2–Q3, regarding each group, were very similar (Table S2).

In 2020, we observed a similar pattern in Q1 to that of 2019. The students presented short answers related to common knowledge, and many of them commented on morphology, ecology and examples (Fig 1B). Few students committed conceptual errors and they were mainly concentrated on morphology and examples (Fig 1D). In Q2, the responses were much more comprehensive, and more students covered topics such as morphology, metabolism, taxonomy, and reproduction (Fig 1B). In turn, fewer students committed conceptual errors, which mainly focused on metabolism (Fig 1D). In Q3, more students commented on morphology, metabolism and reproduction in comparison to Q1 (Fig 1B). As in 2019, the ANOVA suggests that there is a difference between the Shannon indices of the Q1, Q2 and/or Q3 questionnaires (F(2, 88) = 11.586, p < 0.001, η_p_2 = 0.21) (Fig 2B). In general, the richness of the students’ answers increased during the project (H_Q2–Q1_=0.31, 95% CI=[0.18; 0.44], p < 0.001; H_Q3–Q1_=0.35, 95% CI=[0.15; 0.51], p = 0.002) (Fig 2C and Table 1). This increase in richness was maintained for all microorganisms in Q2, with a lower effect size between Q3 and Q1 (Fig 2D and Table 2). However, the confidence intervals are large and, in most cases, the result of the t-test does not discard the possibility of the absence of difference in the Shannon indices between each questionnaire (Table 2). Interestingly, in the virus group, there was a greater increase in richness throughout the project than in the other groups (H_Q2–Q1_=0.48, 95% CI=[0.29; 0.85], p = 0.003; H_Q3–Q1_=0.61 95% CI=[0.25; 0.94], p = 0.024) (Fig 2D and Table 2).

**TABLE 2.** Differences in the Shannon index between the questionnaires Q1 and Q2, and Q1 and Q3 for the 2020 edition.

| **2020 edition** |  |  |  |  |  |  |
| --- | --- | --- | --- | --- | --- | --- |
| **Group** | **Q2-Q1** |  |  | **Q3-Q1** |  |  |
|  | **Effect Size** | **CI 95%** | **p value** | **Effect Size** | **CI 95%** | **p value** |
| **All groups** | 0.31 | [0.18; 0.44] | **<0.001** | 0.35 | [0.15; 0.51] | **0.002** |
| **Archaea** | 0.49 | [0.13; 0.69] | 0.079 | 0.32 | [-0.06; 0.59] | 0.094 |
| **Bacteria** | 0.32 | [-0.09; 1.3] | **<0.001** | 0.29 | [-0.47; 1.12] | 0.596 |
| **Fungi** | 0.29 | [-0.02; 0.46] | 0.323 | 0.17 | [-0.12; 0.37] | 0.426 |
| **Protozoan** | 0.56 | [0.21; 0.92] | 0.010 | 0.38 | [-0.43; 0.74] | 0.867 |
| **Virus** | 0.48 | [0.29; 0.85] | **0.003** | 0.61 | [0.25; 0.94] | 0.024 |
| **Conceptual errors** | -0.20 | [-0.99; 0.31] | 0.503 | -0.30 | [-1.51; 0.19] | 0.056 |

As observed in the 2019 edition, for the 2020 edition, the variation in the richness of the students’ responses between surveys was supported by the calculated Bray-Curtis dissimilarities (Table S3). For 2020, the dissimilarities were slightly greater than in 2019. In general, the dissimilarity was higher than 0.20, and the confidence interval was consistent with values between 0.20 and 0.60 (Table S3). As in the 2019 edition, the main exception was conceptual errors (Q1–Q2=0.68, 95% CI=[0.68; 0.68]; Q2–Q3=0.71, 95% CI=[0.71; 0.71]). Furthermore, the dissimilarity values between Q1–Q2 and Q2–Q3, in terms of each group, were very similar (Table S3), although, in some cases, the point estimate was smaller between Q2–Q3 (as in “All groups”, “Archaea”, and “Fungi”). It is worth noting that in these cases, the 95% confidence intervals remained consistent with similar values in Q1–Q2 and Q2–Q3 (Table S3).

We also used the Bray-Curtis dissimilarities to compare the richness of the students’ responses between the 2019 and 2020 editions (Table 3). Considering each group, the dissimilarity values between 2019 and 2020 in Q1 and Q3 were very similar (Table 3). The exception was the bacteria group, which had a greater dissimilarity in Q3 than in Q1 (Q1: 2019–2020=0.39, 95% CI=[0.30; 0.68]; Q3: 2019–2020=0.64, 95% CI=[0.66;0.86]). Generally, the confidence intervals were wide and included values of dissimilarity between 0.20 and 0.60 (Table 3). Moreover, in most cases, the point estimate dissimilarities were smaller than 0.50. The main exception, as in previous comparisons, were the conceptual errors, which had the largest value of dissimilarity between 2019 and 2020 in all questionnaires (Q1: 2019–2020=0.51, 95% CI=[0.51; 0.51]; Q3: 2019–2020=0.72, 95% CI=[0.72;0.72]).

**TABLE 3.**
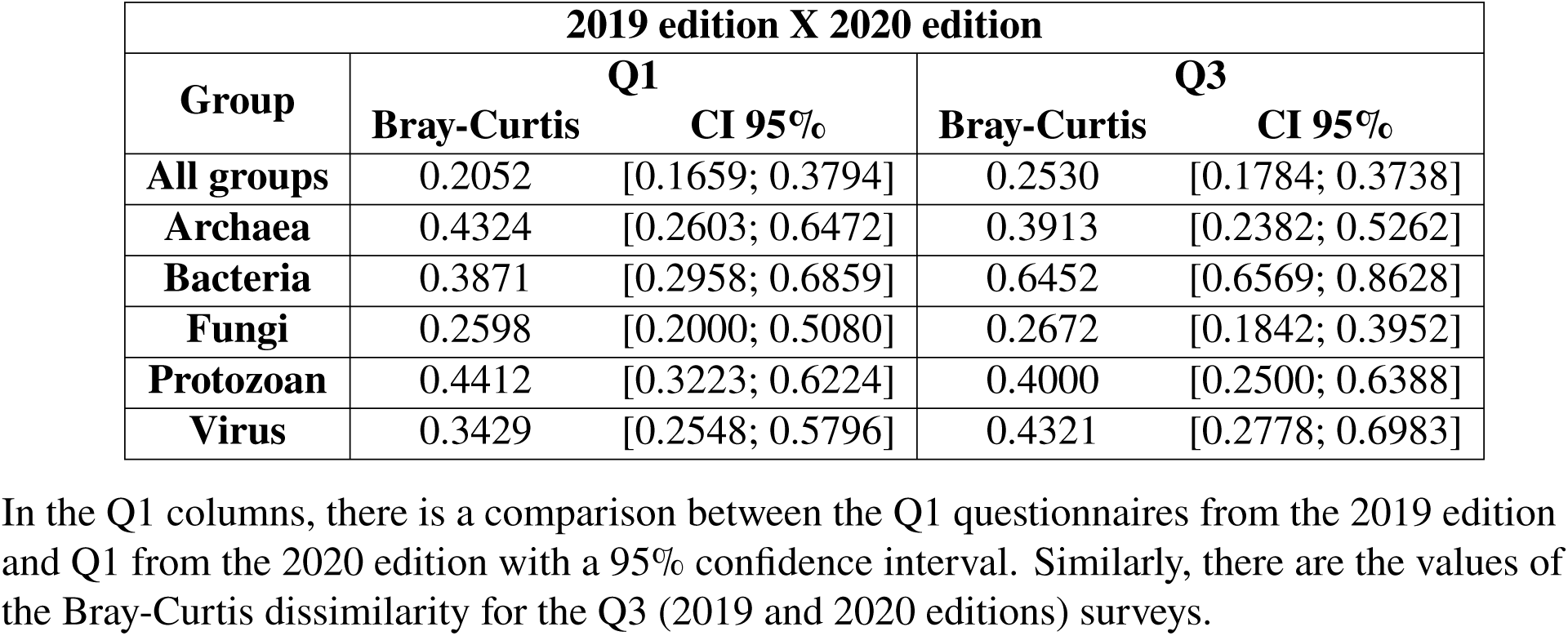
Bray-Curtis dissimilarity of the survey responses between the 2019 and 2020 editions.

| **2019 edition X 2020 edition** |  |  |  |  |
| --- | --- | --- | --- | --- |
| **Group** | **Q1** |  | **Q3** |  |
|  | **Bray-Curtis** | **CI 95%** | **Bray-Curtis** | **CI 95%** |
| **All groups** | 0.2052 | [0.1659; 0.3794] | 0.2530 | [0.1784; 0.3738] |
| **Archaea** | 0.4324 | [0.2603; 0.6472] | 0.3913 | [0.2382; 0.5262] |
| **Bacteria** | 0.3871 | [0.2958; 0.6859] | 0.6452 | [0.6569; 0.8628] |
| **Fungi** | 0.2598 | [0.2000; 0.5080] | 0.2672 | [0.1842; 0.3952] |
| **Protozoan** | 0.4412 | [0.3223; 0.6224] | 0.4000 | [0.2500; 0.6388] |
| **Virus** | 0.3429 | [0.2548; 0.5796] | 0.4321 | [0.2778; 0.6983] |

In the Q1 columns, there is a comparison between the Q1 questionnaires from the 2019 edition and Q1 from the 2020 edition with a 95% confidence interval. Similarly, there are the values of the Bray-Curtis dissimilarity for the Q3 (2019 and 2020 editions) surveys.

According to the “Adopt a Microorganism” evaluation survey, the main positive aspect pointed out by the students in 2019 was the use of Facebook^®^ as a teaching platform due to its accessibility. They also highlighted the autonomy experienced during the learning process, which allowed them to define their study plan and search for the information they found most interesting (Fig 3A). Few negative points were brought out, and they referred mainly to the difficulty of communicating with their group and the ease of losing focus while studying (Fig 3B). In 2020, the students also highlighted the accessibility of Facebook^®^, autonomy, and interaction with the group, as positive aspects (Fig 3C). As in 2019, few negative points were pointed out, and they referred to the difficulty of accessing the Internet, the difficulty of communicating with group members, and distraction by the social network during their research (Fig 3D). Two questions were added to the 2020 questionnaire to assess the importance of the “Adopt a Microorganism” project during the suspension of face-to-face classes due to the COVID-19 pandemic. Most students mentioned that the project allowed them to keep in touch with colleagues during social distancing, mostly because of the routine of publications and challenges posted on Facebook^®^ (Fig 4A). In addition, most of them also pointed out that the project enabled the continuation of learning, even remotely, mainly because it motivated students to research on the proposed themes and encouraged group discussions (Fig 4B).

**Fig. 3.**
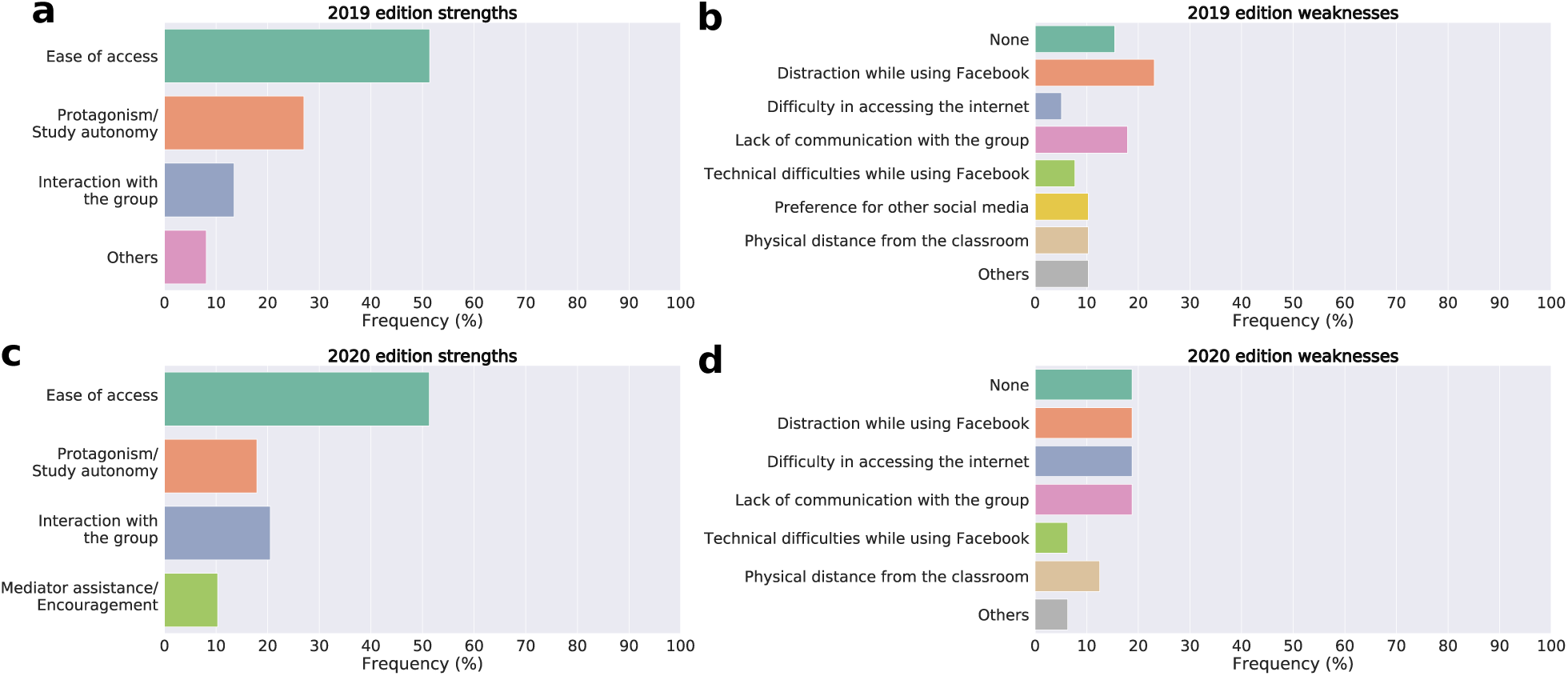
Students’ perceptions of the “Adopt a Microorganism” project. **[a]** positive points highlighted in 2019; **[b]** negative points highlighted in 2019; **[c]** positive points highlighted in 2020; **[d]** negative points highlighted in 2020.

**Fig. 4.**
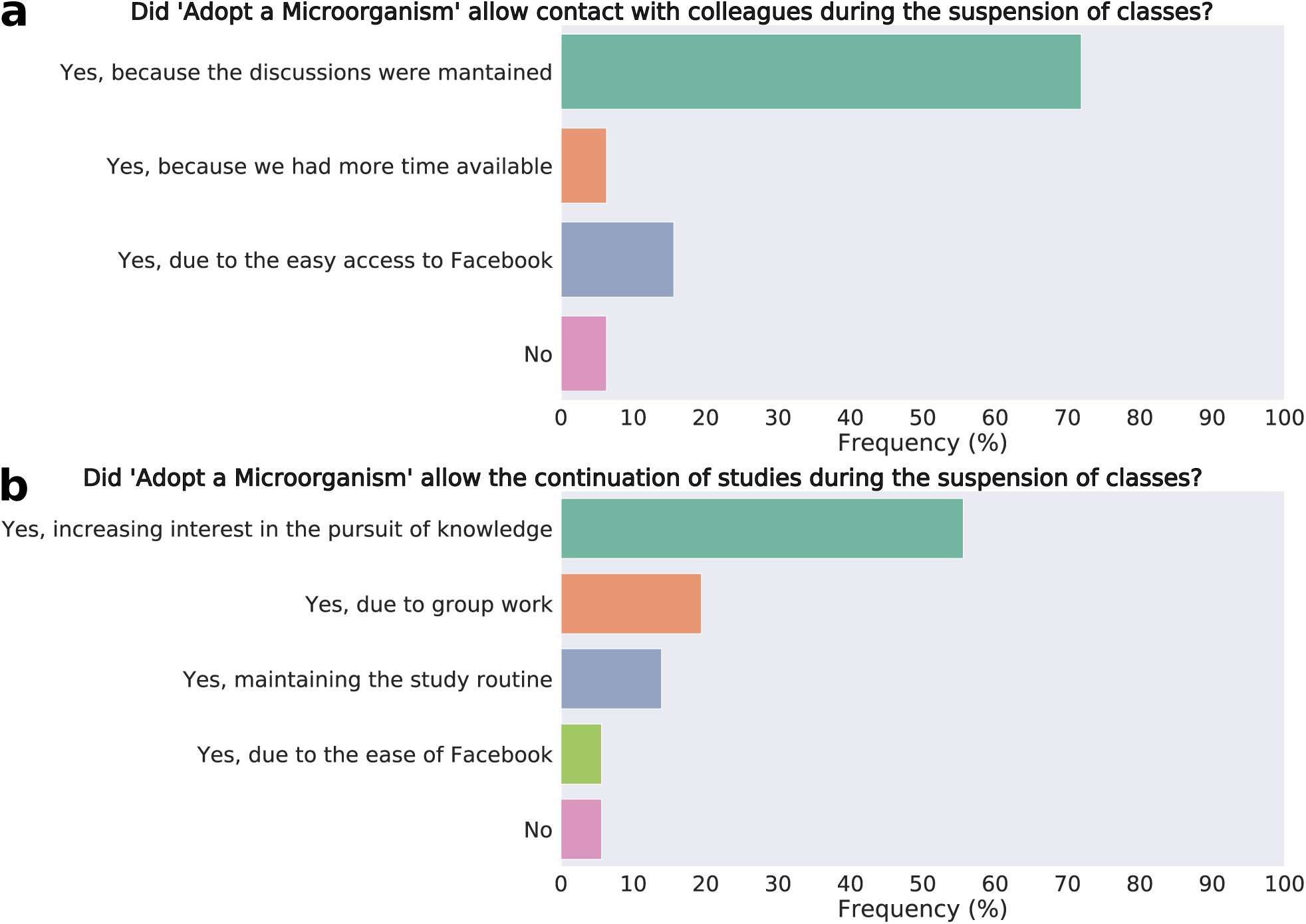
Students’ perceptions regarding the maintenance of “Adopt a Microorganism” during the suspension of face-to-face classes in 2020, due to the COVID-19 pandemic. **[a]** impact of “Adopt a Microorganism” in maintaining contact with colleagues during the pandemic; **[b]** impact of “Adopt a Microorganism” on learning continuity.

## DISCUSSION

The cornerstone of blended learning is the use of the Internet associated with traditional face-to-face classes (28), and this has been the model of the “Adopt a Microorganism” project applied in high school at IFSP, Sorocaba campus. In 2020, however, classes were suspended due to the COVID-19 pandemic, and the project was developed entirely remotely, relying only on the activities and virtual discussions that took place on Facebook^®^. To assess potential differences and similarities between both models, hybrid and remote education, we evaluated the students’ discourse richness throughout the project. An unusual way of analyzing richness in discourse is through the Shannon Diversity Index (16), which is widely used in biology to understand community dynamics and species diversity (18; 17). This index is also commonly used in microbiology, in studies related to microbiome composition (20; 19). In this work, the Shannon index was chosen because the responses to the questionnaires provided us with data on the amount of variety on microbiology subjects, that is, the category diversity (“species”) on a response (“region”). As diverse microbiology vocabulary may facilitate learning about microorganisms (15), we opted to extrapolate this index to analyze the questionnaires and verify potential changes in the content richness of the participating students in both teaching models.

A particular concern with remote learning, especially when applied on an emergency basis, is the decrease in student engagement in the proposed activities (29). In fact, in 2020, we observed a lower commitment in some project activities, especially in the responses to the questionnaires, which were shorter and more objective when compared to the answers obtained in 2019 (Fig. S1). Despite that, the students’ discourse richness was quite similar when comparing the two years (Fig 2). This suggests that, although the information was presented more concisely, remote learning also promoted a greater discourse richness at the end of the project, similar to what we observed in the blended learning format.

In 2020, fewer students committed conceptual errors in comparison to the 2019 edition. This observation may be related to the fact that, in 2020, students were much more concise in their responses. Thus, they would be much less prone to make mistakes. To clarify this, we made a linear regression model (Fig. S3) based on the correlation between the number of words written and the number of mistakes made by the students in their answers. The correlation coefficient suggests that there is a positive correlation between the number of words written and the number of mistakes. Interestingly, the lower regression coefficient in Q2 and Q3 suggests that in the two years, the “Adopt a Microorganism” allowed a reduction in the number of errors by words written by the students. However, it is important to remember that, in 2020, Q2 and Q3 were carried out over the Internet. Therefore, we cannot exclude the possibility that students have consulted materials while filling out these surveys.

During the pandemic period, it has been widely discussed that remote learning decreases student interaction (29; 30), which would be quite worrying since interactions are an essential part of the teaching-learning process (31). In the 2020 edition of “Adopt a Microorganism”, students’ perception of promotion of interaction with colleagues was approximately 75% higher than in the blended edition (Fig 3). Also, approximately 90% of students considered that the “Adopt a Microorganism” allowed them to have contact with their colleagues during class suspension owing to the pandemic (Fig 4A). Jackson (2020) (32) also observed a positive assessment of remote learning and pointed out that students emphasized how much this experience kept them connected. The interaction between students promoted by the “Adopt a Microorganism” is also very likely related to the platform on which it was developed, since social networks, such as Facebook^®^, allow for higher interaction when compared to other platforms used for educational purposes (33). However, we can not attest that this interaction is greater than or equal to what occurred in the blended learning model, but that it existed in both models.

For meaningful learning, there is a mix between verbal and visual resources that allows students to create deeper connections regarding the content, so they retain information more effectively (34; 35). Visual resources were more widely used in “Adopt a Microorganism” during 2019 when the face-to-face hands-on practical activities (“Journey into the World of Microorganisms and Human Parasites”) were held, and the topics covered in those activities were widely represented in the clouds of words of Q2 (Fig. S4), in addition to the huger mention of examples of groups or species of microorganisms (Fig 1A). Since one of the objectives of the “Journey into the World of Microorganisms and Human Parasites” is to introduce students to the “adopted” microorganisms in the form of practical activities (use of microscopes and visits to laboratories), it is possible that this style of activity has increased students’ interest in these themes and facilitated the appropriation of examples.

A remarkable point in carrying out the “Adopt a Microorganism” project in 2020 is the COVID-19 pandemic, which puts viruses at the center of discussions and in the spotlight of all media, directly influencing the retention of concepts related to this microorganism (36). This was evident to us since the students who “adopted” the viruses showed the greatest increase in discourse richness in 2020 (Fig 2D). Also, the difference between the Shannon indices in Q3 and Q1 of the 2020 virus group was the largest compared to the other groups (Table 3). This suggests that students were able to express knowledge about the adopted microorganism even months after the project, which indicates greater knowledge retention. Furthermore, there is also the fact that usually what most catches people’s attention concerning microbiology are viruses and bacteria (36), which, in part, may be related to the data observed in our study.

The use of social networks for educational purposes, such as Facebook^®^, has been widely discussed in the literature, and its benefits and harms are extensively discussed (37; 38; 39; 40). One of the drawbacks is the high distraction capacity these platforms offer, which students highlighted as one of the negative points of the project (Fig 3). In 2019, a small portion of students stated their preference for using other social networks to conduct the “Adopt a Microorganism” (Fig 3B), probably because they had face-to-face classes and other activities that allowed the learned content to be more structured and organized. This preference for other social networks disappeared in 2020. We believe that, because Facebook^®^ was the only formal learning environment available for the students, the platform organization and structure (posting styles and distribution of students in closed groups) may have been interesting and effective leading students to not judge other social networks as more suitable. Students’ perception of the learning environment changes with the current context, and the circumstances imposed by the COVID-19 pandemic may have contributed to a more positive perception of the platform used, precisely because it stimulates the continuity of studies and interpersonal relationships (32).

Another essential point to be considered when applying the “Adopt a Microorganism” is the need for Internet access, an evident problem in the pandemic context, not only in Brazil (41; 42; 43). This was a weakness pointed out by students, especially in 2020, perhaps owing to the impossibility of accessing the school’s computers. In 2020, those who may not have access to the Internet may have found it harder to participate in the project, as well as students who may have had to share their computers with others at home. It is worth mentioning that the students unanimously opted to continue the project after the suspension of classes. Even with possible difficulties in accessing the Internet and without face-to-face lectures, they pointed out that this allowed them to continue their studies during the pandemic (Fig 4B).

In the context of class suspension, encouraging students to continue their studies is very important for the learning process, even though online classes may not be able to engage students and guarantee their motivation (30). However, the students who participated in the “Adopt a Microorganism” stated that the project increased their interest in researching the discussed contents (Fig 4B). This is extremely relevant for us since this is related to one of the main pillars of Education, the “Learning to Learn” (44), which consists of awakening in students an interest in seeking information and promoting their knowledge.

Our results showed that, in both blended and remote learning, students achieved a similar category diversity index and that this occurred regardless of the variety of terms observed in their responses on Q1. Together, these data suggest that the “Adopt a Microorganism” project increases students’ discourse richness in microbiology even without face-to-face traditional classes. Thus, we concluded that, at the end of both models of the “Adopt a Microorganism” methodology, students show a greater discourse richness in their responses, which suggests the acquisition of biology terminology, an essential skill during biology learning.

## Supporting information

Supplementary Figues and Tables

## ACKNOWLEDGMENTS

We thank all the undergraduate students, graduate students and researchers of the Institute of Biomedical Sciences at the University of São Paulo for collaborating as mediators and elaborating the activities. We also thank the high school students of the Federal Institute of São Paulo - Campus Sorocaba for participating in the project.

