## Supplementary Figues and Tables for "Comparison of hybrid learning and remote education in the implementation of the “Adopt a Microorganism” methodology"

### Supplementary Information: "Adopt a Microorganism" methodology enhances the discourse richness of high school students in both hybrid learning and remote education

**Bárbara Rodrigues Cintra Armellini[1,*], Alexandre La Luna[1,2], Vanessa Bueris[1], Alisson Pinto de Almeida[1], Alicia Moraes Tamais[1], Flávio Krzyzanowski Júnior[1,3], Victor Samuel Hasten Reiter[1], Camilo Lellis-Santos[1], and Rita de Cássia Café Ferreira[1]**

[1]Institute of Biomedical Sciences, University of São Paulo, São Paulo, Brazil
[2]Federal Institute of São Paulo, Campus Sorocaba, São Paulo, Brazil
[3]Federal Institute of São Paulo, Campus Capital, São Paulo, Brazil
[4]Federal University of São Paulo, Campus Diadema, São Paulo, Brazil
[*]

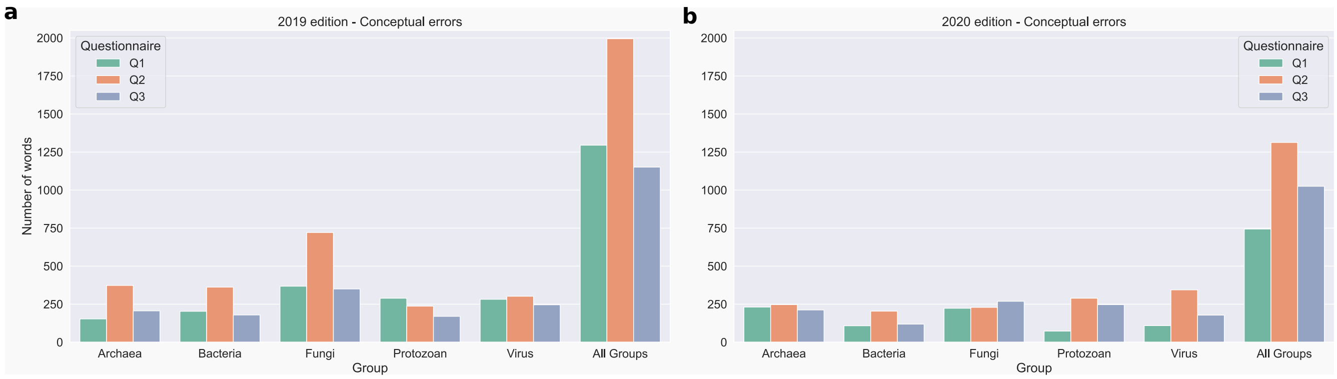

**Figure S1.** Number of words in students' answers. **[a]** 2019 edition; **[b]** 2020 edition.

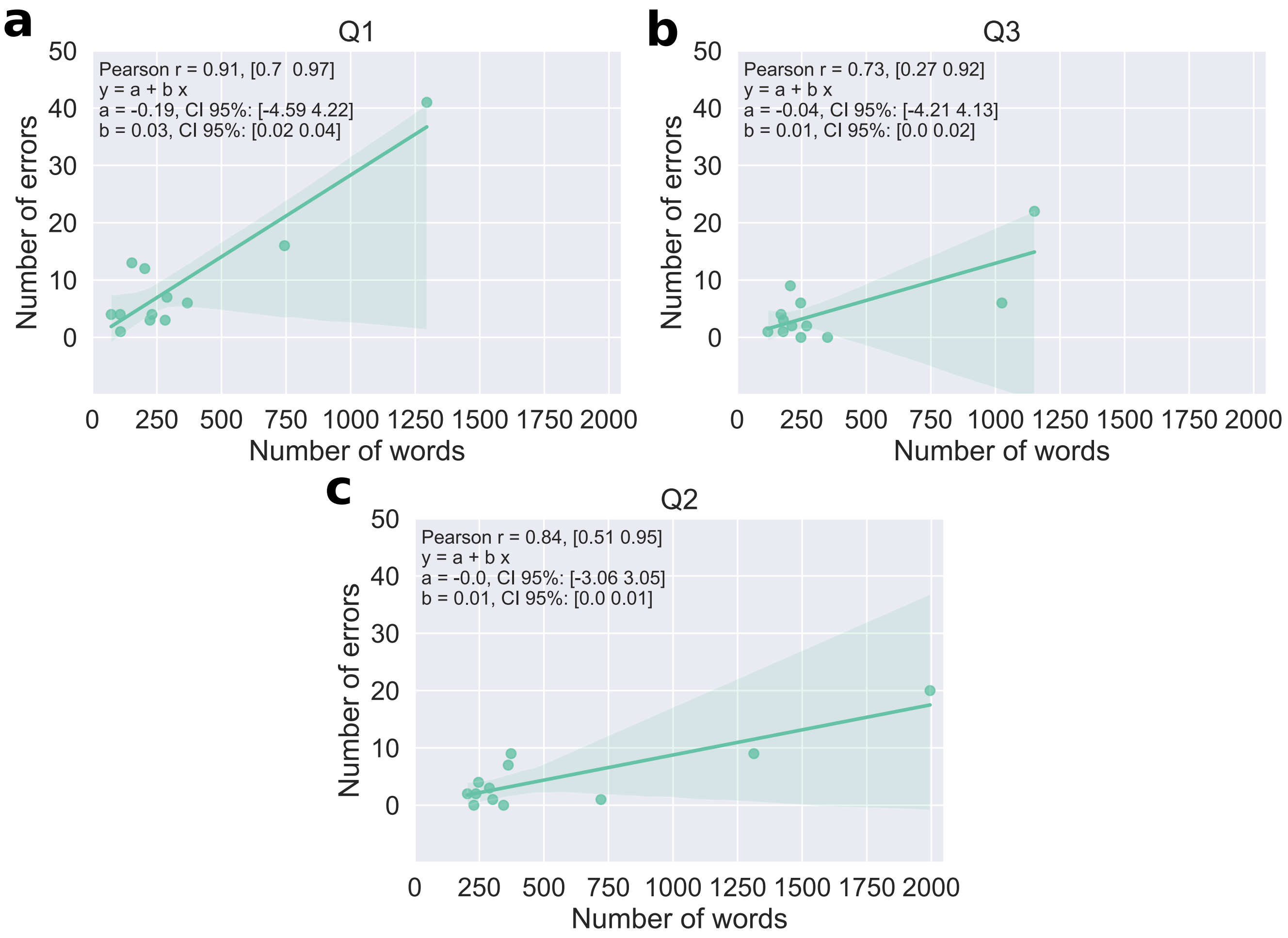

**Figure S2.** Correlation between number of words and number of errors in all answers in 2019 and 2020 edition. In each plot is indicated the Pearson's correlation coefficient and its confidence interval; as well as the linear regression model, with the intercept (a) and slope (b) terms and their respective confidence interval . **[a]** Q1 questionnaire; **[b]** Q2 questionnaire and **[c]** Q3 questionnaire.

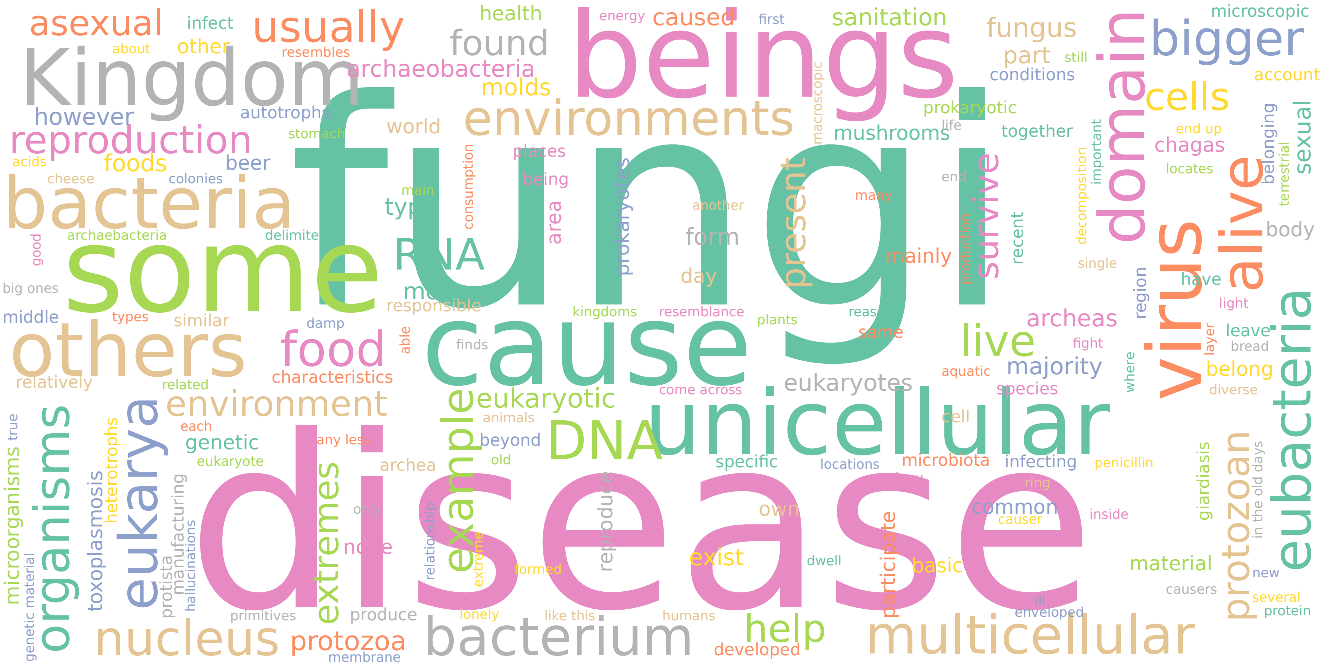

**Figure S3.** Word cloud of the answers in Q2 questionnaire during 2019 edition.

**Table S1.** Bray-Curtis dissimilarity calculated with the survey responses of the 2019 edition.

| 2019 edition | | | | |
|---|---|---|---|---|
| **Group** | **Q1-Q2** | | **Q2-Q3** | |
| | **Bray-Curtis** | **CI 95%** | **Bray-Curtis** | **CI 95%** |
| **All groups** | 0.20 | [0.18; 0.35] | 0.20 | [0.16; 0.32] |
| **Archaea** | 0.41 | [0.31; 0.71] | 0.39 | [0.31; 0.53] |
| **Bacteria** | 0.20 | [0.14; 0.40] | 0.37 | [0.28; 0.47] |
| **Fungi** | 0.34 | [0.28; 0.48] | 0.30 | [0.24; 0.43] |
| **Protozoan** | 0.19 | [0.14; 0.38] | 0.33 | [0.22; 0.45] |
| **Virus** | 0.52 | [0.45; 0.71] | 0.44 | [0.40; 0.61] |

**Table S2.** Bray-Curtis dissimilarity calculated with the survey responses of the 2020 edition.

| 2020 edition | | | | |
|---|---|---|---|---|
| **Group** | **Q1-Q2** | | **Q2-Q3** | |
| | **Bray-Curtis** | **CI 95%** | **Bray-Curtis** | **CI 95%** |
| **All groups** | 0.30 | [0.28; 0.51] | 0.14 | [0.16; 0.40] |
| **Archaea** | 0.44 | [0.33; 0.58] | 0.27 | [0.16; 0.53] |
| **Bacteria** | 0.43 | [0.42; 0.81] | 0.47 | [0.48; 0.89] |
| **Fungi** | 0.30 | [0.24; 0.58] | 0.19 | [0.18; 0.45] |
| **Protozoan** | 0.53 | [0.52; 0.73] | 0.42 | [0.27; 0.58] |
| **Virus** | 0.50 | [0.42; 0.74] | 0.40 | [0.24; 0.59] |

**Table S3.** Difference in the Shannon index between the questionnaires Q1 and Q2, and Q1 and Q3 for 2019 edition.

| 2019 edition | | | | |
|---|---|---|---|---|
| **Group** | **Q2-Q1** | | **Q3-Q1** | |
| | **Effect Size** | **CI 95%** | **Effect Size** | **CI 95%** |
| **All groups** | 0.17 | [0.07; 0.29] | 0.08 | [-0.05; 0.18] |
| **Archaea** | 0.21 | [-0.18; 0.80] | 0.11 | [-0.14; 0.72] |
| **Bacteria** | 0.27 | [0.06; 0.62] | 0.16 | [0.01; 0.52] |
| **Fungi** | 0.74 | [0.55; 0.95] | 0.19 | [-0.03; 0.47] |
| **Protozoan** | -0.2 | [-0.41; 0.05] | -0.13 | [-0.77; 0.12] |
| **Virus** | -0.21 | [-0.49; 0.08] | -0.11 | [-0.29; 0.18] |

**Table S4.** Difference in the Shannon index between the questionnaires Q1 and Q2, and Q1 and Q3 for 2020 edition.

| **2020 edition** | | | | |
|---|---|---|---|---|
| **Group** | **Q2-Q1** | | **Q3-Q1** | |
| | **Effect Size** | **CI 95%** | **Effect Size** | **CI 95%** |
| **All groups** | 0.31 | [0.18; 0.44] | 0.35 | [0.15; 0.51] |
| **Archaea** | 0.49 | [0.13; 0.69] | 0.32 | [-0.06; 0.59] |
| **Bacteria** | 0.32 | [-0.09; 1.3] | 0.29 | [-0.47; 1.12] |
| **Fungi** | 0.29 | [-0.02; 0.46] | 0.17 | [-0.12; 0.37] |
| **Protozoan** | 0.56 | [0.21; 0.92] | 0.38 | [-0.43; 0.74] |
| **Virus** | 0.48 | [0.29; 0.85] | 0.61 | [0.25; 0.94] |
