## Supplementary material for "Comparison of hybrid learning and remote education in the implementation of the “Adopt a Microorganism” methodology": S2 Appendix

### ADOPT A MICROORGANISM PROJECT EVALUATION QUESTIONNAIRE

**1-** Please indicate, according to your opinion, the positive points in using Facebook® as a tool to learn about microorganisms.

**2-** Now indicate the negative points in the use of Facebook®.

**3-** How do you evaluate the participation of the mediators?

**4-** How could mediators improve their participation?

**5-** Below are some statements (A to D) related to the use of Facebook® in the learning process. Evaluate how much each of these statements contributed to your learning about Biology in the Adopt a Microorganism project. For this evaluation, use the following scale:

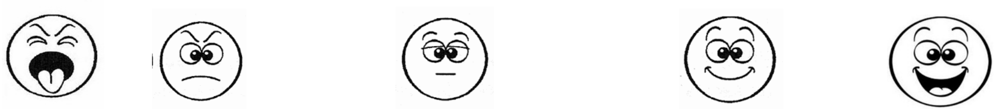

| Terrible Bad | Bad | More or Less | Good | Excellent |
|---|---|---|---|---|

A- I was able to learn and study when and where I wanted.

B- I had more contact with classmates, teachers and the mediators to clarify doubts about the subject.

C- I was able to participate more actively in my own learning.

D- Easier to ask questions than in the classroom.

**6-** We're done! Would you like to comment on anything else? Be my guest!
