## Supplementary material for "Comparison of hybrid learning and remote education in the implementation of the “Adopt a Microorganism” methodology": S1 Appendix

### RULES - ADOPT A MICROORGANISM ON FACEBOOK

- Each group must make at least ONE post and THREE comments per challenge.
- The challenges will only be validated if they are accompanied by a personal comment on the topic and bring a new information/question about the micro-organism. The mediators will decide if the challenges are valid and will indicate that by commenting "Validated" on each of the comments and posts made by the group.
- The participation grade for this activity will be achieved if the minimum number of posts/shares and comments are made.
- To conduct information research you should use reliable sources, whether formal (scientific articles, for example) or informal (such as newspapers, magazines, science blogs, and others). It is important that these sources are identified in the group posts.

#### 1 - GENERAL CHALLENGES

Two giant viruses are discovered in Brazil
Fábio de Castro, O Estado de S.Paulo
27 February 2018

Two new giant viruses have been discovered in Brazil, according to a study published Tuesday, 27, in the journal Nature Communications. The two specimens - which belong to a new genus baptized Tupanvirus - have a genetic complexity never found in any other virus, according to the authors of the study.
These virus are so large that they can be observed under a common optical microscope, but they do not cause disease and preferentially infect amoebas, according to one of the authors of the study, Jonathan Abraham, professor at the Department of Microbiology at the Federal University of Minas Gerais (UFMG). Unlike other viruses, Tupanvirus has a kind of tail, whose function is still unknown.
"Like other giant viruses discovered in the past, Tupanvirus infects amoebas. The difference is that it is much more generalist: unlike the others, it is able to infect different types of amoebas,", said Abrahão.

According to him, amoebas are among the oldest beings on Earth, which leads scientists to hypothesize that giant viruses may also be quite old. "Looking at the relationship between giant viruses and amoebas is equivalent to looking into the past and understanding the origin of the first life forms,", explained the scientist.

##### Challenge 1

The text cites the three current domains of life: Archaea, Bacteria, Eukarya. These domains comprise different cell types. Compare the three domains by considering the presence or absence of a nucleus, types of cell envelope, presence or absence of organelles, ability to reproduce, and list the main differences between them.

**Challenge 2**
The text describes all the organisms we know today. Considering the definition of "Life"proposed by NASA - "self-sustaining chemical system capable of Darwinian evolution", name which of the domains presented in the text, plus viruses, would be considered alive according to NASA's definition.

**Challenge 3**
Still commenting on the diversity of organisms listed in the text, consider now the definition proposed by other scientists: "For something to be considered alive, it must have metabolism (a set of chemical reactions that a being performs to obtain energy)". Evaluate this definition considering the domains already mentioned and viruses.

#### 2 - SPECIFIC CHALLENGES

##### 2.1 - Fungi
**Challenge 1**

What is the largest living being?

Strange World - Superinteressante Magazine

Published Apr 18, 2011

The largest creature on the planet was only discovered in 1996: a fungus growing under the ground in the Malheur National Forest in Oregon. This *Armillaria ostoyae*, popularly known as the "honey mushroom", was born as a tiny particle, impossible to see with the naked eye, and extended its filaments over an estimated period of 2,400 years. From the surface, you can see only its ends along the tree trunks, but underground it covers 880 hectares - the equivalent of 1,220 soccer fields. "It still grows from 70 centimeters to 1.20 meters a year," says the agronomist João Lúcio de Azevedo, from USP. Before its discovery, the largest living being was another fungus of the same species, found in 1992. Until the 1990s, the title belonged to a sequoia tree in California.

Question

Did you know that until 1969 the honey mushroom and the sequoia belonged to the same kingdom? Name some characteristics of plants and fungi taking into account the morphology (shape), the cell structures, and the means of obtaining and storing energy.

**Challenge 2**
Some fungi are unicellular (yeast) and others are multicellular (molds and mushrooms). Comment on the main macroscopic and microscopic differences between yeast and mold. How is the reproduction in these two organisms?

**Challenge 3**

Have you ever heard of "sick building syndrome"? Hint: it has to do with new building forms and fungi!

Research about this syndrome and describe what type of fungus it is related to and what characteristics facilitate its spread in these environments.

**2.2 - Viruses**
**Challenge 1**
Viruses are small obligate intracellular parasites that have no metabolism of their own. They use the host cell to synthesize their components, thus allowing their perpetuation in nature. They can contain DNA or RNA as genetic material. Some viruses have a layer called the “envelope" (eg. dengue viruses), which consists of a lipid-rich membrane that envelops the viral particle externally. On the other hand, viruses that do not have this layer are said to be non-enveloped (eg. HPV viruses). In addition to the classification by their structures, viruses can be allocated into groups according to their propagation characteristics in nature. For example, arboviruses are the designation given to viruses that are transmitted by insects to vertebrate hosts. These viruses have a very wide geographical distribution spanning all continents, in both temperate and tropical regions.

Question

Compare the structures of some viruses (I-Dengue; II-Zika; III-Yellow fever; IV-Chikungunya) according to the following characteristics:

a) Presence or absence of envelope.
b) Type of genetic material (RNA or DNA).
c) Insects capable of transmission to humans or animals.

**Challenge 2**

What are STIs?

Sexually Transmitted Infections (STI) are caused by viruses, bacteria or other microorganisms. They are transmitted mainly through sexual contact (oral, vaginal, anal) without the use of a male or female condom, with a person who is infected. The transmission of an STI can also occur from mother to child during pregnancy, delivery or breastfeeding. The treatment of people with STIs improves their quality of life and interrupts the chain of transmission of these infections. The care and treatment are free of charge in the SUS - Health Public System.

The terminology Sexually Transmitted Infections (STI) is now being adopted in place of Sexually Transmitted Diseases (STD), because it highlights the possibility of a person having and transmitting an infection, even without signs and symptoms.

Question

HPV infection is one of the most prevalent diseases in the world. Women, men and even children are susceptible to HPV. It is estimated that at least half of all sexually active people have had contact with the virus.
Can you tell which are the diseases caused by HPV? If half of the sexually active population has had contact with the virus, why don't all of them develop the diseases caused by it?

**Challenge 3**

Previously we talked about STIs (Sexually Transmitted Infections) such as those caused by HPV and HIV. Since 2014, the Brazilian Ministry of Health has offered the quadrivalent vaccine that protects against four types of HPV.

Question: Is there a vaccine against HIV? What are the difficulties to produce it?

##### 2.3 - Protozoa

**Challenge 1**

Among the diseases that occur in untreated sewage and water, Giardiasis stands out. It is caused by the protozoan Giardia intestinalis, which causes symptoms such as diarrhea and vomiting, and consequently affects the child's nutrition. If this condition occurs frequently in the first five years of life, the sequelae can be limiting and definitive.

Question

Considering the text above, explain the relationship between poor sanitation and the risk of acquiring Giardiasis. Is it possible to acquire the disease even in developed, clean cities? Name some ways.

**Challenge 2**

Toxoplasmosis is an infectious disease caused by *Toxoplasma gondii*, an intracellular parasitic protozoan that can infect birds, rodents, wild animals, and a large number of mammals (cattle, pigs, goats, sheep), including humans of all ages.

What is the reason why toxoplasmosis is known as the "cat disease"? What are the characteristics of Toxoplasma? What is its life cycle like?

**Challenge 3**

In September 2017, in the city of Marilândia (ES), there was an outbreak of Toxoplasmosis in a daycare center, where 18 children and two female employees were diagnosed with the disease.

At the time of the outbreak, the city's health secretary stated that she did not know the cause of the contagion, but that she was already trying to find out.
Source: Gazeta Online

Question:

Can you come up with at least two hypotheses to determine where people got infected in the daycare center?

**2.4 - Archaeobacteria**

**Challenge 1**

Yellowstone National Park is one of the most visited national parks in the United States. This park is home to the famous Great Prismatic Spring, which has specific environmental conditions, such as low pH, high temperature (70 ºC) and high concentration of minerals.
In 2016, a young man and his girlfriend were visiting the park and decided to take a bath in a thermal lake. As they approached the lake to test the temperature, the young man fell into the water and died. The body was not found, possibly because the high temperatures and acidity of the water dissolved his tissues. However, even in these extreme conditions, certain organisms thrive, such as Archaea and bacteria, which are responsible for the vibrant colors of these bodies of water.

Since 1969, living things have been grouped into five kingdoms: Animalia, Plantae, Protist, Monera, and Fungi, as proposed by researcher Robert Whittaker. In 1977, researchers Carl Woese and George Fox proposed the organization of all living things into 3 domains: Bacteria, Archaea and Eukarya. This model was based on shared structural features, use of coenzymes, and the similarities and differences between nucleotide sequences of ribosomal RNA (rRNA).

Question
Explain why the Archaea have been classified into a single domain. Is this group closer to Bacteria or Eukarya?

**Challenge 2**

Have you ever heard of a primitive earth? Is it possible to make any connection between the predominant features of this period and the different metabolism of the Archaea?

**Challenge 3**

Do you remember the Big Spring in Yellowstone Park where the temperature reached 70ºC? In your opinion, what factors (DNA, proteins, structures and cellular mechanisms) would allow Archaea to survive in such extreme environments?

#### 2.5 Eubacteria

**Challenge 1**

Outbreak of *Escherichia coli* infection in Canada and the United States due to contamination of vegetables.

Canada - The Public Health Agency of Canada, along with the Food Inspection Agency and Health Canada, have investigated an outbreak of *Escherichia coli* type O157. The outbreak involved five eastern provinces. Individuals became ill in November and early December 2017. Based on the results of the investigation during the outbreak, exposure to lettuce was identified as the source of the outbreak, but the cause of the contamination was not identified. No individuals had the illness after December 12, 2017. United States - According to the U.S. Centers for Disease Control and Prevention (CDC), an *Escherichia coli* outbreak caused 25 people to become ill between November 15 and December 8. Two of the hospitalized patients developed hemolytic uremic syndrome, a type of kidney failure. Although the source of the infection is still unknown, the CDC is investigating leafy greens and romaine lettuce.
The species *Escherichia coli* stands out for their importance in clinical medicine and biotechnology. Strains of this bacterial species are important constituents of the intestinal microbiota of animals and are therefore considered fecal coliform.

Question

How might food in general be contaminated with Escherichia coli? What are safe food handling procedures?

**Challenge 2**

What are the possible ways that bacteria reproduce? Which one is used by Escherichia coli?

**Challenge 3**

The bacterium *Escherichia coli* is widely used in current biotechnology processes. Can you tell which characteristics allow this kind of manipulation? And how is it used in these processes?
